# Whole Genomes Define Concordance of Matched Primary, Xenograft, and Organoid Models of Pancreas Cancer

**DOI:** 10.1101/209692

**Authors:** Deena M.A. Gendoo, Robert E. Denroche, Amy Zhang, Nikolina Radulovich, Gun Ho Jang, Mathieu Lemire, Sandra Fischer, Dianne Chadwick, Ilinca M. Lungu, Emin Ibrahimov, Ping-Jiang Cao, Lincoln D. Stein, Julie M. Wilson, John M.S. Bartlett, Ming-Sound Tsao, Neesha Dhani, David Hedley, Steven Gallinger, Benjamin Haibe-Kains

**Affiliations:** Princess Margaret Cancer Centre, University Health Network, Toronto, Ontario, Canada; PanCuRx Translational Research Initiative, Ontario Institute of Cancer Research (OICR), Toronto, Ontario, Canada; Informatics and Bio-computing Program, Ontario Institute for Cancer Research, Toronto, Ontario, Canada; Princess Margaret Living Biobank Core, University Health Network, Toronto, Ontario, Canada; Department of Statistical Science, University of Toronto, Toronto, Ontario, Canada; Department of Pathology, University Health Network, University of Toronto, Toronto, ON, Canada.; UHN Program in BioSpecimen Sciences, Department of Pathology, University Health Network, Toronto, Ontario M5G 2C4, Canada.; Transformative Pathology, Ontario Institute for Cancer Research, Toronto, Ontario, Canada; Division of Medical Oncology, Princess Margaret Cancer Centre, Toronto, Ontario, Canada; Molecular Genetics Department, University of Toronto, Toronto, Ontario, Canada; Lunenfeld-Tanenbaum Research Institute, Mount Sinai Hospital, Toronto, Ontario, Canada; Hepatobiliary/Pancreatic Surgical Oncology Program, University Health Network, Toronto, Ontario, Canada; Department of Medical Biophysics, University of Toronto, Toronto, Ontario, Canada; Department of Computer Science, University of Toronto, Toronto, Ontario, Canada

**Keywords:** Disease models, Xenograft, Organoid, Bioinformatics, Genomics, Whole-Genome Sequencing, PDAC, Pancreatic Cancer, Cancer

## Abstract

Pancreatic ductal adenocarcinoma (PDAC) has the worst prognosis among solid malignancies and improved therapeutic strategies are needed to improve outcomes. Patient-derived xenografts (PDX) and patient-derived organoids (PDO) serve as promising tools to identify new drugs with therapeutic potential in PDAC. For these preclinical disease models to be effective, they should both recapitulate the molecular heterogeneity of PDAC and validate patient-specific therapeutic sensitivities. To date however, deep characterization of PDAC PDX and PDO models and comparison with matched human tumour remains largely unaddressed at the whole genome level. We conducted a comprehensive assessment of the genetic landscape of 16 whole-genome pairs of tumours and matched PDX, from primary PDAC and liver metastasis, including a unique cohort of 5 ‘trios’ of matched primary tumour, PDX, and PDO. We developed a new pipeline to score concordance between PDAC models and their paired human tumours for genomic events, including mutations, structural variations, and copy number variations. Comparison of genomic events in the tumours and matched disease models displayed single-gene concordance across major PDAC driver genes, and genome-wide similarities of copy number changes. Genome-wide and chromosome-centric analysis of structural variation (SV) events revealed high variability across tumours and disease models, but also highlighted previously unrecognized concordance across chromosomes that demonstrate clustered SV events. Our approach and results demonstrate that PDX and PDO recapitulate PDAC tumourigenesis with respect to simple somatic mutations and copy number changes, and capture major SV events that are found in both resected and metastatic tumours.

## INTRODUCTION

Pancreatic ductal adenocarcinoma (PDAC) is a highly lethal, therapy-resistant malignancy, with a dismal overall 5-year survival rate that remains minimally unchanged over the past several decades **[1, 2]**. Multiple failed clinical trials suggest that new approaches are necessary towards understanding PDAC molecular etiology and personalizing treatment **[3]**. There is continued interest in *in vitro* and *in vivo* preclinical models that emulate the PDAC morphologic and genomic landscape, and which can ultimately serve as platforms to select and test candidate treatments.

An increasing number of experimental findings demonstrate that patient-derived organoids (PDO) and patient-derived xenografts (PDX) function as important preclinical platforms for investigations into the molecular landscape of cancer. Studies on cell lines and PDX have alluded to the agreement of tumour histo-architecture between disease models and primary human PDAC **[4, 5]**. Huang *et al*. demonstrated that PDAC PDO maintain differentiation status, recreate histo-architectural heterogeneity, and retain patient-specific physiological changes **[6]**. Recent studies emphasized the fidelity of PDAC disease models at the genomic level by focusing on mutational profiles from whole-exome sequencing (WES) data. Xie *et al* **[7]**, characterized somatic SNVs (singlenucleotide variations) of paired primary tumours and metastases and PDX, focusing on the distribution of allelic frequencies and functional mutations affecting known cancer drivers or tumour suppressors. Witkiewicz et al. and Knudsen et al. **[4, 8]** compared cell lines and PDX models derived from the same tumour, demonstrating their utility in recapitulating patient-specific therapeutic sensitivities. Collectively, these studies provide valuable insight on the significance of PDAC models as ‘avatars’ for precision treatment, but their singular focus on mutational patterns and morphological changes fails to capture the full spectrum of complex genomic events that underlie PDAC heterogeneity.

Despite progress in sequencing efforts for PDAC, comprehensive assessment of PDAC disease models using whole-genome sequencing (WGS) has not been performed. Using WGS, the genomic complexity of resected PDAC tumours has been thoroughly described **[2, 9–12]**. WGS analysis of primary and metastatic tumours has also shed light on catastrophic mitotic phenomena, such as chromothripsis, that occur with high frequency in the disease **[12]**. WGS analysis of PDAC preclinical models would demonstrate how such models recapitulate complex genomic events, including structural variation (SV) and copy number variation (CNV) changes that play a significant role in PDAC tumourigenesis and drug response **[13–15]**.

A large majority of PDAC disease model literature has focused on cell lines and PDX, while genomic characterization of PDO models remains unaddressed. This is despite growing findings that suggest that PDO, compared to PDX and cell lines, may present as models that can reconstitute niches most similar to PDAC **[6, 16]**. In particular, the 3-D architecture of organoids promotes interaction between pancreatic cells (including normal pancreatic cells, paraneoplastic cells, and neoplastic pancreatic cells) and provides improved conditions for polarization of these epithelial cells **[17, 18]**. Organoids have also been shown to exhibit ductal- and stage-specific characteristics, and recapitulate the full spectrum of PDAC tumourigenesis **[16]**. These promising findings pose an opportunity for probing such PDOs at the genomic level. Genomic analysis of PDOs remains a missing link to identify whether these models recapitulate patient tumours at the molecular level, a necessary step before widespread screening therapeutics.

In this study, we conducted a detailed assessment of the genetic landscape of a series of paired tumours and PDX from primary PDAC and liver metastases, including WGS data from a unique set of 5 matched ‘trios’ of primary tumour, PDX, and PDO. Our analysis demonstrates that PDXs and PDOs succeed in recapitulating genome-wide copy number changes of their donor tumours, and mutational events across major PDAC driver genes. While structural variation events showed high variability at genome-wide and chromosome-centric levels, we highlight striking cases of donor-model concordance across chromosomes with large clustering of SV events. Our results provide new insights into the interplay and fidelity of different disease models toward recapitulating PDAC heterogeneity and genomics.

## RESULTS

We conducted comprehensive characterization of paired PDX and PDO from PDAC primary tumours and liver metastases [Figure 1, Supplemental_Table_S1.pdf, Supplemental_Fig_S1.pdf]. WGS was performed for 16 pairs of tumours with their matched disease models [Supplemental_Table_S1.pdf, Supplemental_Fig_S1.pdf]. These included a series of 10 resected tumours and 6 liver metastasis [Supplemental_Table_S1.pdf]. Of the primary tumours, five samples had matching PDO, comprising a unique cohort of matched tumour-PDX-PDO, hereby referred to as ‘trios’ throughout the paper. The PDOs were derived from the PDX tissue, as opposed to the primary patient material.

**Figure 1:**
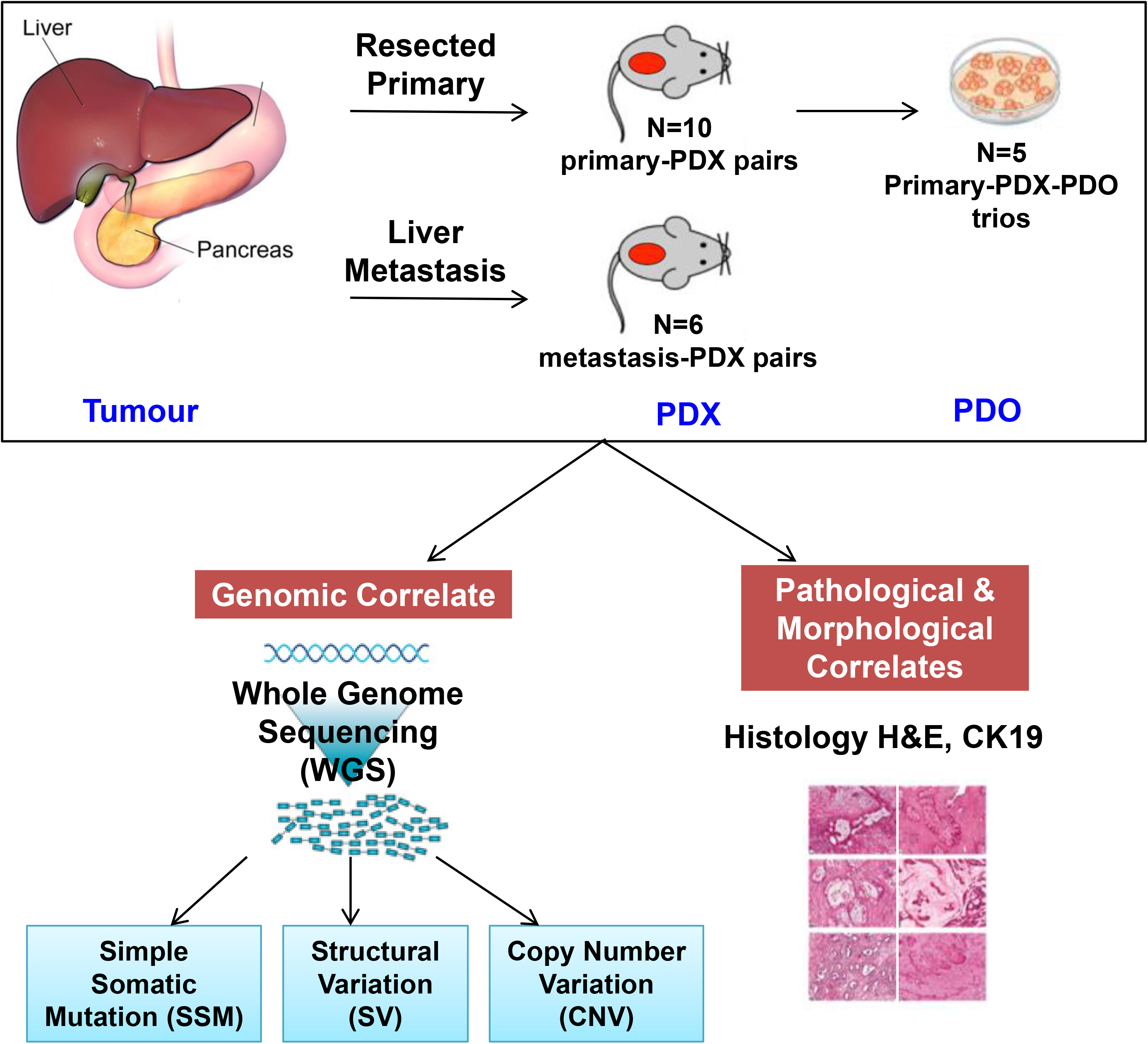
Schematic overview of samples and analysis. Disease models (PDX and PDO) were compared against matched human tumours in terms of morphological agreement, and genomic agreement. Assessment of genomic agreement was established by a top-down approach that determined genomic changes at varying levels of complexity, spanning single-based resolution (SSM) towards genome-wide comparisons (CNV).

### Histology

We assessed the conservation of tumour histo-architecture in the trios [Figure 2]. All samples had strong staining of cytokeratin 19 (CK19) [Supplemental_Fig_S2.pdf, Supplemental_Table_S2.pdf], confirming that all PDX and PDO samples consisted of human PDAC. All samples demonstrated a mainly tubular architecture with varying degrees of cellularity and stroma content [Supplemental_Table_S2.pdf]. PDOs were also observed to mimic the histo-architectural heterogeneity of their matched tumours, and were comprised of a hollow central lumen surrounded by one layer of polarized epithelia [Figure 2, Supplemental_Fig_S2.pdf].

**Figure 2:**
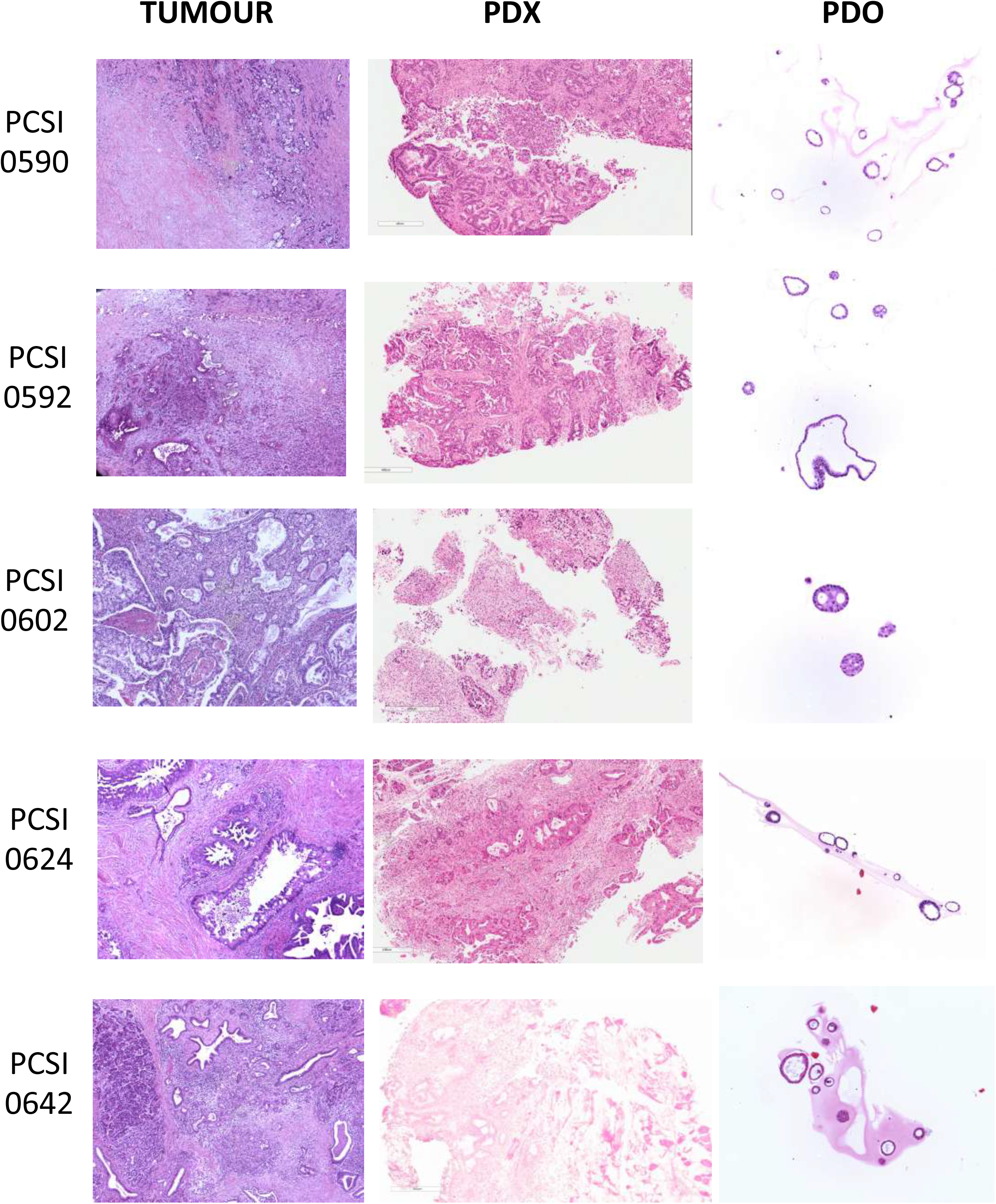
Representative H&E stained sections of primary tumour, matched PDX, and PDO in 5 trios. PDX images are shown at 5.6X zoom, with a scale of 400 uM. PDO images are shown at 10X zoom.

### Genomic profile - SSM, SV, and CNV - of PDAC driver genes

We compared the genomic profiles of 9 PDAC driver genes, including oncogenes and tumour suppressor genes, in primaries and metastases, and in their matched disease models [Figure 3]. We annotated SSM, SV breakpoints, and copy number changes associated with these genes in tumour-PDX pairs, and in tumour-PDX-PDO trios. Several structural arrangements, including deletion events and copy number loss that were observed in the tumours, were recapitulated in the corresponding PDX (eg: MAP2K4 across primary-PDX pairs) [Figure 3]. Homozygous deletions of CKDN2A were observed in several primary-PDX pairs (ex: PCSI_0590, PCSI_0642) and recapitulated in the trios [Figure 3].

**Figure 3:**
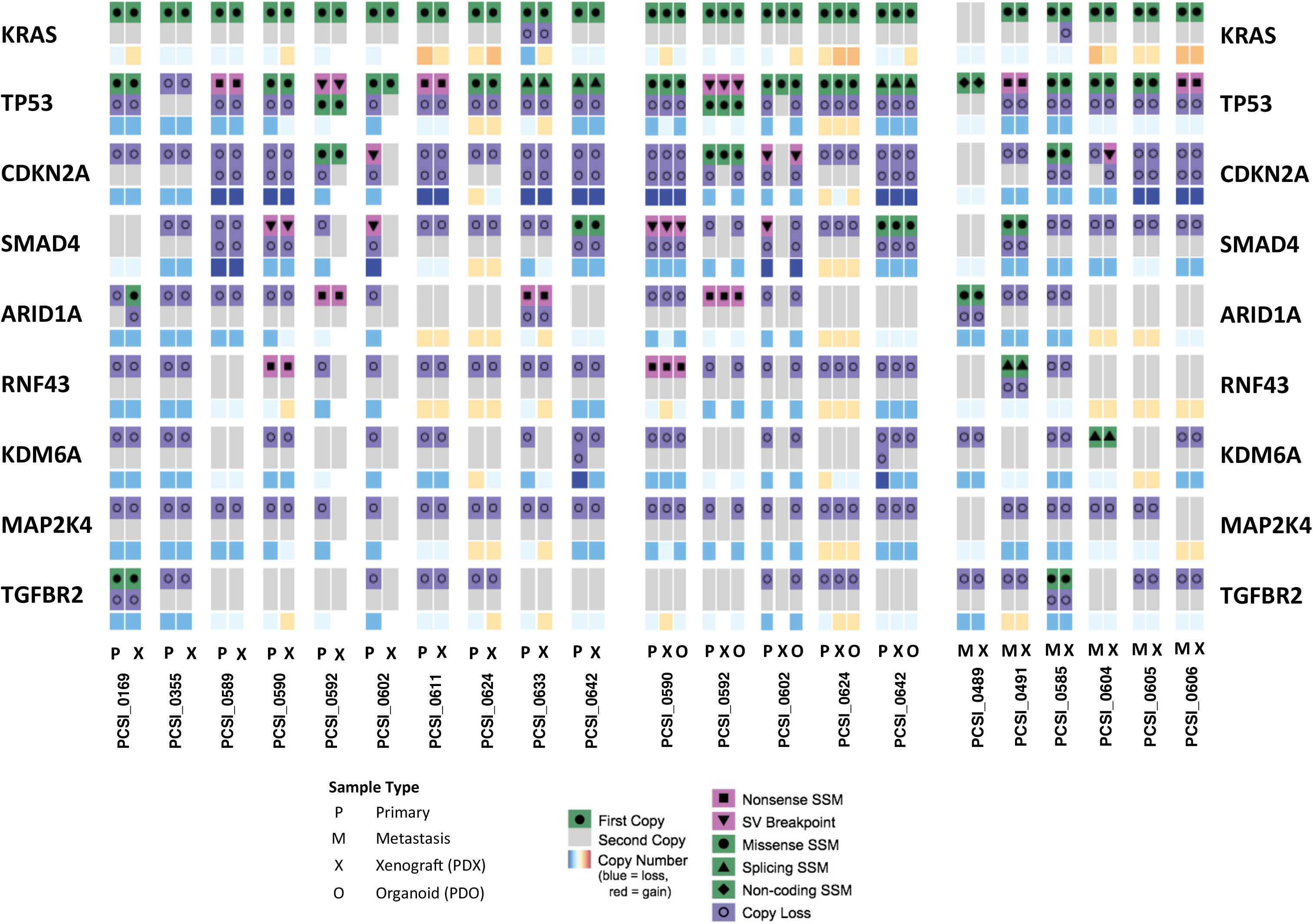
Comparative analysis of SSM, SV, and CNV across genes of PDAC tumourigenesis and disease therapy. The genomic profile of driver genes across primary-PDX pairs (left), metastasis-PDX pairs (middle), and primary-PDX-PDO trios (right) is shown.

### Simple Somatic Mutation

We conducted a comparative analysis of SSM events in tumour-PDX primaries, trios, and paired tumour-PDX metastases, to assess preservation of somatic variants between disease models and their source tumours [Figure 4A, Supplemental_Table_S3.pdf (A-E)]. An average of 4,500 and 5,000 mutations were identified in the resected tumours and in their matched PDX, respectively [Figure 4A, Supplemental_Table_S3.pdf (A)]. More than 50% of the mutations observed in the primary tumour were retained in the paired PDX[Figure 4A]. To assess conservation of synonymous and non-synonymous mutational events (SNVs) in the pairs, we calculated the Jaccard index for each tumour-PDX pair for 12 mutation types observed in the samples [Figure 4B]. Among all mutation types, 7/10 tumour-PDX pairs demonstrated strong concordance of mutation calls (Jaccard index ≥ 0.6) [Figure 4B Supplemental_Table_S3.pdf (D)], with PCSI_0355 as the almost perfect concordant pair in this series (Jaccard index = 0.8). The remaining 3/10 pairs (PCSI_0169, PCSI_0589, PCSI_0602) had moderate concordance (Jaccard index 0.51 to 0.57). Functional mutation types (missense, nonsense) were strongly concordant in >60% of the pairs (Jaccard index ≥ 0.6). Pairwise-comparison of mutation categories also highlighted other mutation types, including mutations of lincRNA, for which the majority of pairs (>60% of the pairs) exhibited strong concordance (Jaccard index ≥ 0.6).

**Figure 4:**
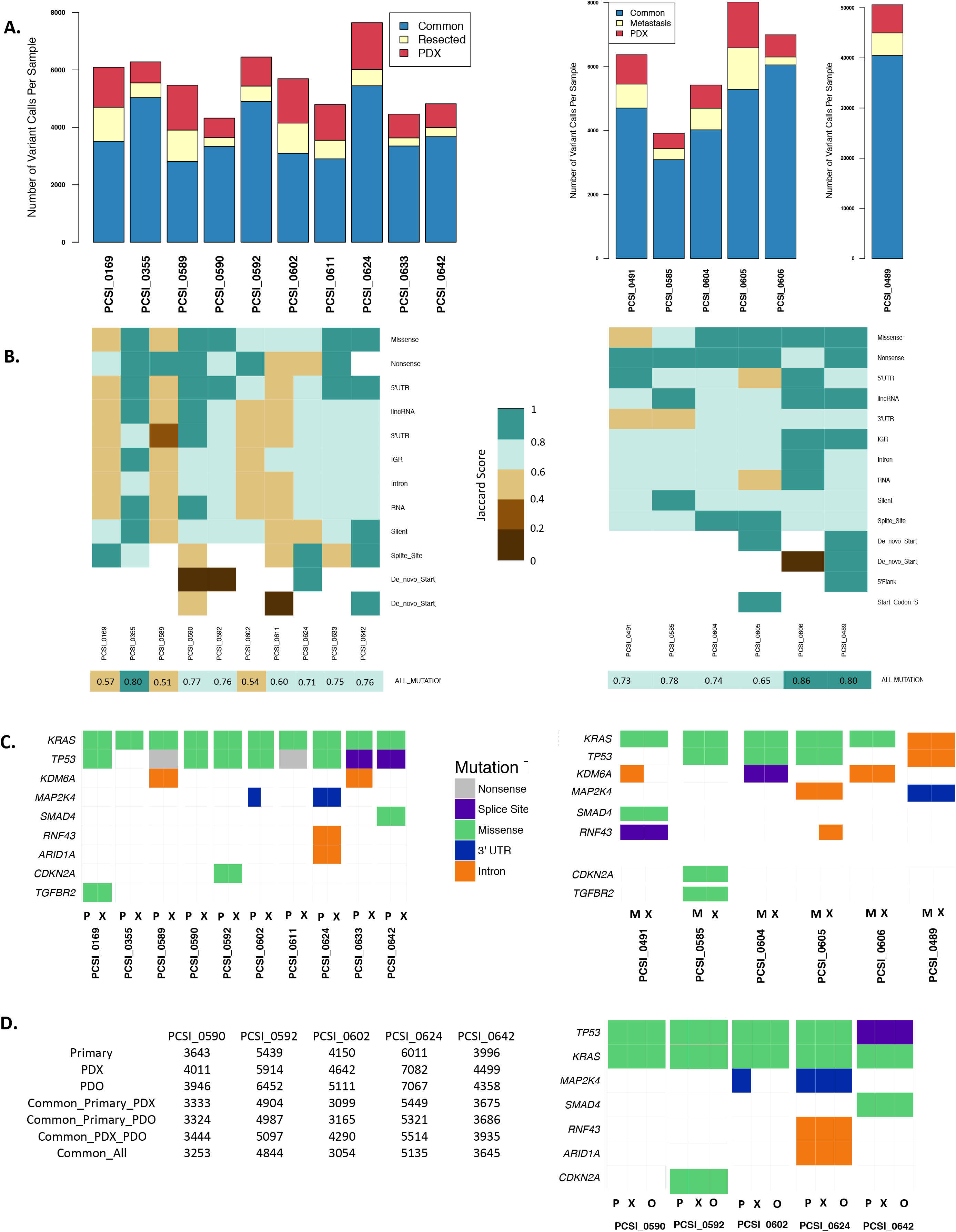
Comparative analysis of SSM across tumour-PDX pairs of resected primary (left) and liver metastasis (right). **(A)** Total number of variant calls across matched tumour-PDX pairs for 10 primary samples (left) and 6 metastasis samples (right). The total number of common mutations across a given pair is indicated, as well as variants that are specific to the tumour sample or matching PDX. **(B)** Heatmap representation of the Jaccard index for a given tumour-PDX pair, across all categories of functional and non-functional mutation types annotated in the primary samples (left) and metastasis samples (right). White cells indicate mutation types that are not available for a tumour-PDX pair. Overall concordance of a tumour-PDX pair across all mutations is indicated by the Jaccard index in the last row (“ALL_MUTATIONS”). **(C)** Conservation of mutation types across oncogene and tumour suppressor genes in the primary samples (left) and metastases (right). Samples are labeled as primary (P), xenograft (X), and metastasis (M). **(D)** Total number of SSM calls across primary-PDX-PDO trios. The top rows indicate the total number of mutations observed in each for the primary, PDX, and PDO samples. Common mutations across primary-PDX, primary-PDO, and PDX-PDO pairs also indicated. The total number of common mutations shared across all samples of the trio is delineated in the last row. Conservation of mutation types across oncogenes and tumour suppressor genes in the trios is also indicated (right). Samples are labeled as primary (P), xenograft (X), and organoid (O).

We conducted an in-depth analysis of mutation patterns for driver genes of PDAC tumourigenesis, including PDAC oncogenes (KRAS) and tumour suppressor genes (CDKN2A, TP53, SMAD4, TGFB2) [Figure 4C]. In cases of apparent discordance between tumour-PDX pairs for a given mutation, we conducted additional manual inspection of the variants to ensure that the failure to call the variant was not attributed to insufficient coverage or statistical threshold for variant calling. Despite discrepancies observed between tumour-PDX pairs across different mutation types, the main driver genes remained highly conserved. KRAS and TP53 mutations were most frequently observed (9/10 pairs) and harbored the same functional consequences (mostly missense mutations) in matched tumour-PDX samples [Figure 4C]. Detailed examination of the KRAS mutations calls in the primary tumour series revealed the presence of G12D, G12R, and G12V oncogenic mutations, with the majority of missense mutations belonging to G12D. The majority of TP53 mutations were missense mutations, but also included splice-site mutations in 2 pairs of the series [Figure 4C]. We assessed the frequency of reads carrying the variant alleles across tumour-PDX pairs. Comparable frequencies between tumours and matched PDX were also observed in larger sets of variants from genes encompassing oncogenes, tumour suppressor genes, and frequently mutated genes involved in PDAC tumourigenesis [Supplemental_Fig_S3.pdf].

Comparison of liver metastasis pairs [Figure 4B] recapitulated much of the somatic mutational landscape observed in the primary tumours. Notably, there was a higher average of mutational calls in the 5 liver metastasis samples (~5,300 mutations per tumour) and their matched PDX (~5,500 mutations per PDX), with a range of 66–86% overlap of mutation calls in metastasis-PDX pairs [Figure 4A, Supplemental_Table_S3.pdf (B)]. One sample (PCSI_0489) had a strikingly high mutation load, with an average of ~45,500 mutations in both the metastasis sample and the matched PDX, due to DNA mismatch-repair (MMR) deficiency. This sample had a germline frameshift MLH1 deletion (not shown), loss of heterozygosity of MLH1 on the p-arm of chromosome 3 and elevated C>T transitions, which corresponds with the diagnosis of Lynch syndrome in the patient **[19]**. All of the metastasis-PDX pairs, including the MMR deficient case, exhibited moderate to almost perfect concordance of mutations in all mutational categories (Jaccard index across ‘ALL_MUTATIONS’> 0.6) [Figure 4B, Supplemental_Table_S3.pdf (E)]. The PCSI_0606 pair was almost perfectly concordant (Jaccard index = 0.86). Missense mutations for KRAS and TP53 were observed in the majority of the metastasis-PDX pairs; for the MMR case only nonfunctional variants for those genes were observed [Figure 4C]. Frequencies of reads carrying the variant allele were also comparable between liver metastasis samples and their matched PDX [Supplemental_Fig_S4.pdf].

Matched PDO samples demonstrated the same mutation pattern in oncogenic driver and tumour suppressor genes as that of their matched PDX and tumour [Figure 4D, Supplemental_Table_S3.pdf (C)]. Pairwise comparisons across the tumours and the models highlighted overall consistency of read frequencies carrying the variant alleles in tumour-PDX, tumour-PDO, and PDX-PDO samples for each of the trios [Supplemental_Fig_S5.pdf].

### Structural Variation

We used our WGS data to assess structural variation (chromosomal rearrangements). Analysis of SVs in the resected primary-PDX pairs revealed that the majority of variants were intra-chromosomal, including deletions (DEL), inversions (INV), and duplications (DUP) [Figure 5A, Supplemental_Table_S4.pdf (A)]. Inter-chromosomal translocations (TRA) were less prevalent [Figure 5A, Supplemental_Table_S4.pdf (A)]. The total numbers of SV events observed were similar between primary-PDX pairs with the exception of PCSI_0611, where the number of events in the PDX were almost double that observed in the primary tumour [Figure 5A].

**Figure 5:**
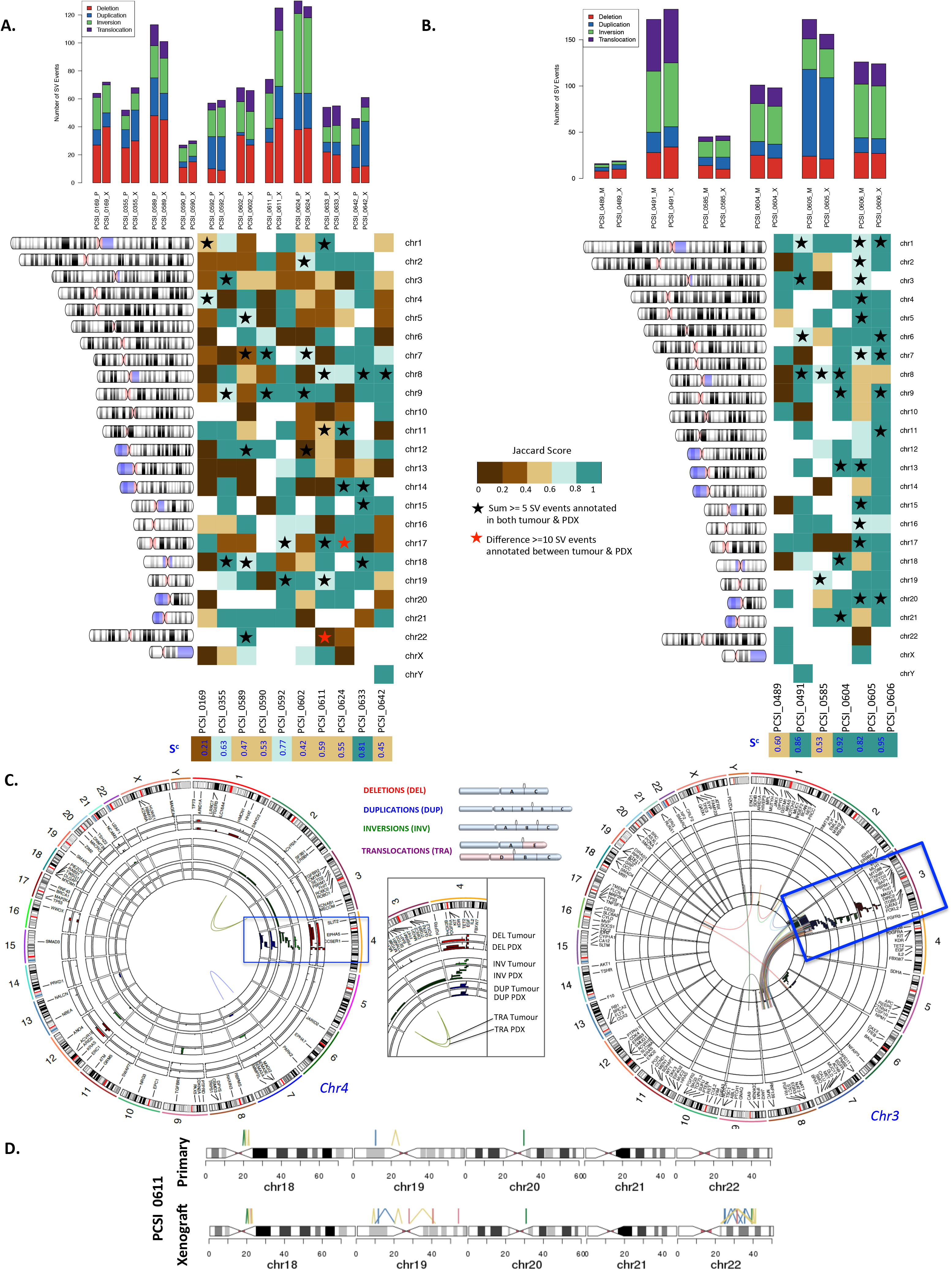
Comparative analysis of structural variation (SV) across tumour-PDX pairs of the primary and metastasis cohorts. **(A)** Distribution of SV events (deletion, duplication, inversion, translocation) in each sample across 10 primary-PDX pairs (left) and 6 metastasis-PDX pairs (right). **(B)** Chromosome-specific Jaccard indices across 10 primary-PDX pairs (left) and 6 metastasis-PDX pairs (right). Samples are labeled by their PCSI identifier. Chromosomes with an observed large number of rearrangements (≥5 events) in both the tumour and matched PDX are indicated (black stars). Chromosomes that are discordant between a tumour-PDX pair (≥10 SV events difference) are highlighted (red stars). The overall concordance (S^c^) score for a tumour-PDX pair across all chromosomes in indicated (bottom row). **(C)** Genome-wide SV events observed in tumour-PDX pairs in PCSI_0169 (resected primary, right) and PCSI_0491 (liver metastasis, left). Each type of SV event is color-coded with a similar color between tumours and matching PDX. For each SV type, tumours are annotated on the outer rings of the circos plot and the matching PDX on the inner rings. Chromosomes exhibiting clustered SV events (potential chromothripsis) are highlighted in the blue box. **(D)** Comparison of SV events across chr18-chr22 for the PCSI_0611 primary tumour and its matching PDX. There is an apparent chromothripsis event on chr22 of the PDX but not the primary sample.

We compared the distribution of structural variation events across each chromosome, for every primary-PDX pair [Figure 5B, Supplemental_Table_S5.pdf (A)]. Strong (Jaccard index ≥ 0.6) and almost perfect (Jaccard index ≥ 0.8) concordance was observed across many chromosomes with elevated numbers of SV events (defined as ≥5 SV events in both the primary and its matching PDX). This concordance extended to chromosomes that displayed clusters of structural variants in the primary-PDX pair. Elevated counts of clustered chromosomal rearrangements in these chromosomes, compared to the rest of the genome, is suggestive of chromothripsis [Figure 5C, Supplemental_Table_S6.pdf (A)]. In the majority of the primary-PDX pairs, we identified particular chromosomes with clustered SV events, including PCSI_0169 (chr 4), PCSI_0589 (chr 7), PCSI_0592 (chr 17, chr 18), PCSI_0611 (chr1), and PCSI_0642 (chr 8) [Figure 5B, Figure 5C, Supplemental_Fig_S6.pdf, Supplemental_Table_S5.pdf (A)]. We also identified chromosomes that were strongly discordant, owing to a large difference in the number of SV events between the primary and matched PDX. Most notable of these was the q arm of chromosome 22 in PCSI_0611, for which a cluster of SV events was observed in the PDX but not in the primary sample [Figure 5D]. We computed an overall concordance (S^c^) score for each primary-PDX pair, to quantify concordance across all chromosomes that harbor SV events [Figure 5B]. Few of the primary-PDX pairs demonstrate concordance across the genome (S^c^ ≥ 0.6), with the exception of PCSI_0633 (S^c^= 0.81). PCSI_0169 was the least concordant across the genome (S^c^= 0.21). Upon further investigation, we found that the PDX has more than twice as many indels as the primary tumour, and is enriched for deletions, which suggests that mutations may be accumulating due to deficiency in a DNA repair pathway.

We conducted a comparative analysis of SV events in paired metastasis and matched PDX samples [Figure 5] to identify distinct chromosomal instability profiles observed in primary tumours that could also extend to liver metastasis pairs. Metastasis-PDX pairs exhibited variable counts of SV events, ranging between 16-172 events in the samples [Figure 5B, Supplemental_Table_S4.pdf (B)]. Not surprisingly, the MMR deficient case (PCSI_0489) had the fewest number of SV events in the pairs [Figure 5B, Supplemental_Table_S4.pdf (B)], and scored moderately in overall concordance (S^c^ = 0.6). In contrast to the primary-PDX pairs however, the majority of the metastasis-PDX pairs (4/6) demonstrated almost perfect concordance across the genome (S^c^ < 0.8 for PCSI_0491, PCSI_0604, PCSI_0604, PCSI_0606). Structural variation events were either strongly concordant or almost perfectly concordant across the majority of chromosomes in the metastasis-PDX pairs (Jaccard index ≥ 0.6) [Supplemental_Table_S6.pdf (B)], including chromosomes with elevated counts of SV events in both the metastasis and matched PDX [Figure 5B, Supplemental_Table_S5.pdf (B)]. None of the metastasis-PDX pairs harbored discordant chromosomes [Figure 5B]. We identified clustering of SV events in specific chromosomes for the metastasis-PDX pairs, with the exception of the MMR deficient case [Figure 5C, Supplemental_Fig_S7.pdf]. These chromosomes include PCSI_0491 (chr 3), PCSI_0604 (chr 13), PCSI_0605 (chr1, chr2), and PCSI_0606 (chr1, chr9, chr11) [Figure 5C, Supplemental_Fig_S7.pdf, Supplemental_Table_S5.pdf (B), Supplemental_Table_S6.pdf (B)].

Analysis of the primary-PDX-PDO trios highlighted the extent to which structural variation of primary-PDX pairs were captured in organoid models [Figure 6, Supplemental_Fig_S8.pdf]. The distribution of intra-chromosomal and inter-chromosomal events in the PDO samples shares a similar pattern to that of their matched tumour and PDX [Figure 6A, Supplemental_Table_S4.pdf (C)]. By splitting the trio into matched pairs (primary-PDX, primary-PDO, and PDX-PDO) we assessed concordance in the primary tumours and each of the disease models (tumour-PDX, tumour-PDO pairs), and the interrelationship between the disease models themselves (PDX-PDO pairs) [Figure 6B, Supplemental_Table_S5.pdf (C), Supplemental_Table_S6.pdf (C)]. Tumour-PDX and tumour-PDO scores ranged from strong discordance to moderate (S^c^ <0.6) in the five trios [Figure 6B, Supplemental_Table_S6.pdf (C)]. However, strong concordance (Jaccard index ≥ 0.6) and almost perfect concordance (Jaccard index ≥ 0.8) was observed for chromosomes that demonstrate ≥ 5 SV events, including for example, PCSI_0592 (chr17, chr19). This concordance was similarity extended to PDX-PDO pairs in PCSI_0592 (chr17, chr19), PCSI_0602 (chr7, chr9), PCSI_0624 (chr11, chr14), and PCSI_0642 (chr8) [Figure 6B, Supplemental_Table_S6.pdf (C)], suggesting a strong agreement in the trios for chromosomes with clustered SV events. For the tumour-PDX pairs with highly clustered SV events, clustering of structural variation events was also recapitulated in the PDOs (e.g., PCSI_0642, chromosome 8) [Figure 6D, Supplemental_Fig_S8.pdf].

**Figure 6:**
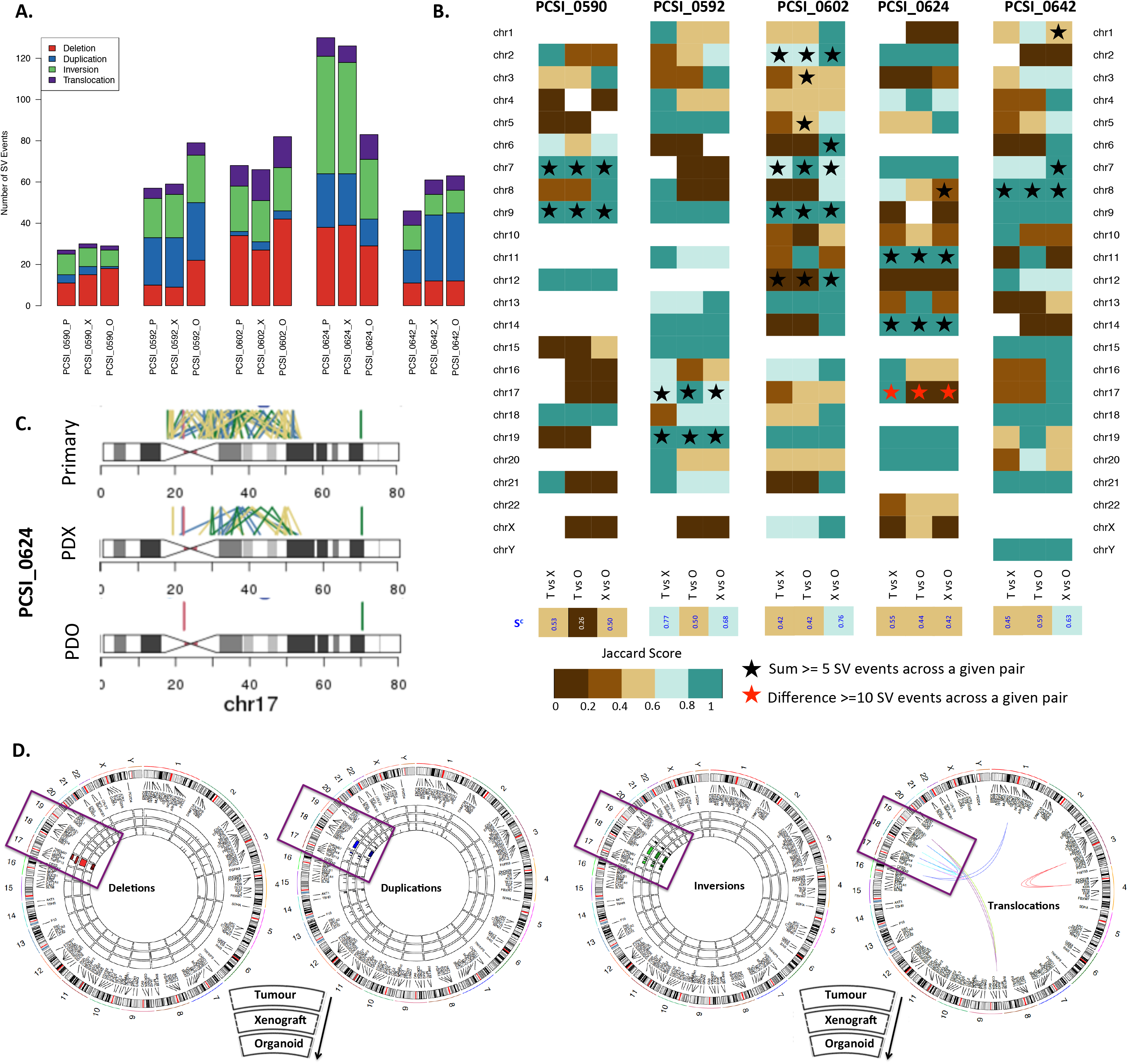
Comparative analysis of structural variation (SV) across tumour-PDX-PDO trios. **(A)** Distribution of SV events (deletion, duplication, inversion, translocation) in 5 trio samples. The primary (P), PDX (X), and PDO (O) sample for each trio is indicated. **B)** Chromosome-specific Jaccard indices in 5 primary-PDX-PDO trios. Each trio is split into 3 pairs representing primary-PDX (T vs X), primary-PDO (T vs O) and PDX-PDO (X vs O) comparisons. Chromosomes with an observed large number of rearrangements (≥5 events) in each pair are indicated (black stars). Chromosomes that are discordant members of a pair (≥10 SV events difference) are highlighted (red stars). The overall concordance (S^c^) score for each pair across all chromosomes in indicated (bottom row). **(C)** Comparison of SV events across chromosome 17 of PCSI_0624. This chromosome was discordant between primary-PDX, primary-PDO, and PDX-PDO pairs. **(D)** Distribution of SV events across the primary tumour, matched PDX, and matched PDO samples of the PCSI_0592 trio. Each type of SV (deletion, inversion, duplication, and translocation) is represented as one circos plot, with 3 rings indicating tumour (outer), PDX (middle), and PDO (inner). Chromosomes demonstrating chromothripsis are highlighted in boxes.

Comparison of tumour-PDX, tumour-PDO, and PDX-PDO pairs highlighted particular cases that were discordant at both chromosome and genome-wide levels. In the trios, we observed the most discordance in all specimens of the trio for chromosome 17 of PCSI_0624. Further investigation into this chromosome revealed a very high count of SV events in the primary tumour, with reduced events in the matching PDX, and almost no events in the PDO [Figure 6C, Supplemental_Table_S5.pdf (C)]. Comparison of the trios genome-wide (using overall S^c^ scores) also demonstrates that PCSI_0590 and PCSI _0624 were the lowest scoring among the trios; all of the pairs of those trios ranged from moderate discordance to weak (S^c^ < 0.6).

### Copy Number Variation

Genome-wide copy number state was assessed for primary-PDX pairs, trios, and metastasis-PDX pairs [Figure 7, Supplemental_Table_S7.pdf]. Evaluation of ploidy could not be determined unambiguously for two of the samples (PCSI_0592 and PCSI_0602) and they were excluded from the copy number analysis. Accordingly, 8/10 primary-PDX pairs were further evaluated for ploidy and copy number state [Supplemental_Table_S7.pdf (A), Supplemental_Fig_S9.pdf]. Both tumours and their matched PDX exhibited comparable ploidy in the majority of the pairs [Figure 7A, Supplemental_Table_S7.pdf (A)]. Two pairs (PCSI_0590 and PCSI_0633) demonstrated a doubling in ploidy in the PDX, compared to the matching primary tumour [Figure 7A, Supplemental_Fig_S10.pdf].

**Figure 7:**
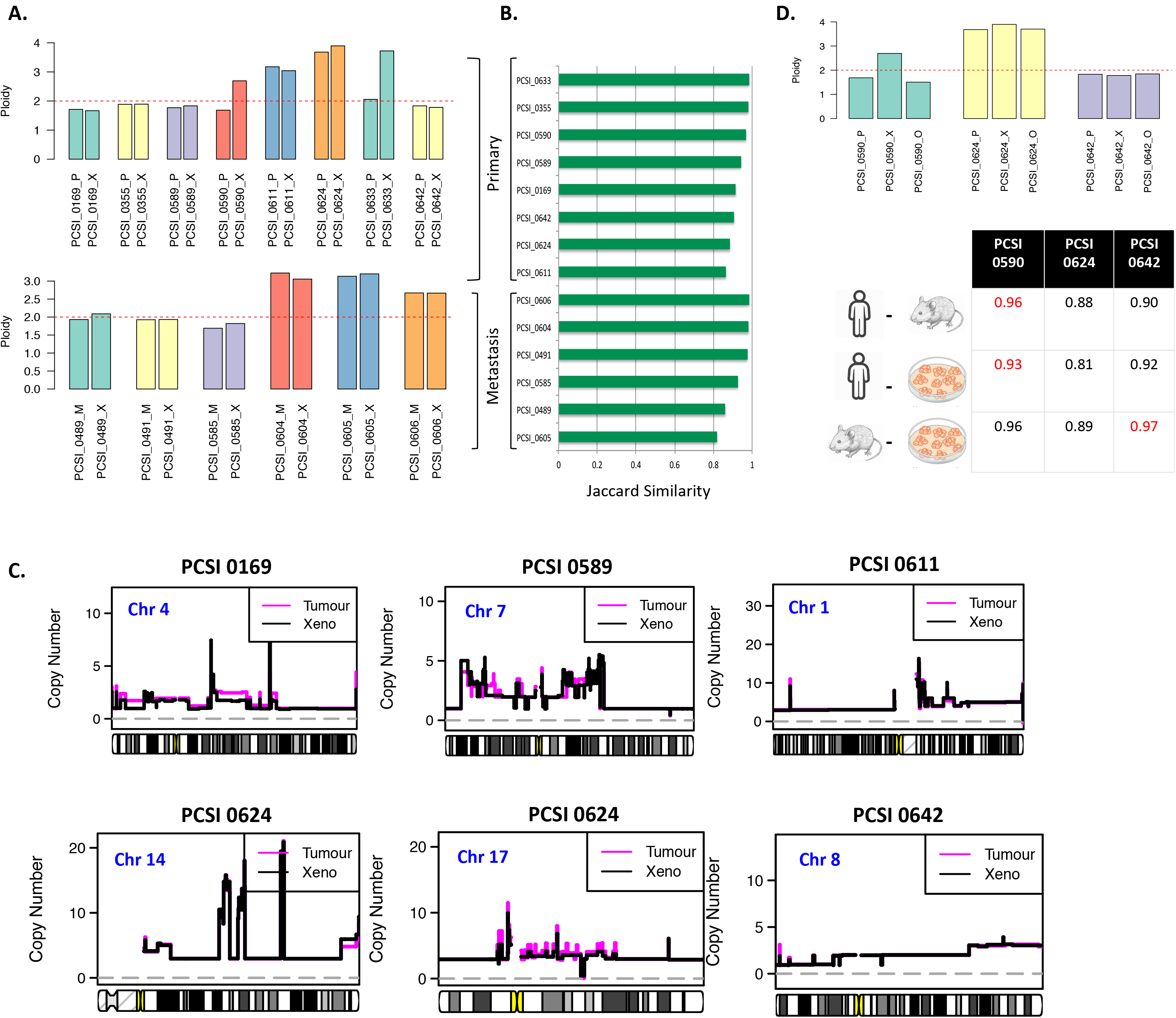
Comparative analysis of copy number state across tumour-PDX pairs of the resected primary cohort. **(A)** Ploidy across tumour-PDX pairs for primary tumours (top) and liver metastasis (bottom). **(B)** Genome-wide concordance score for a given tumour-PDX pair. **(C)** Copy number state for tumour (magenta) and matching PDX (black), highlighted for chromosomes of the primary-PDX pairs for which chromothripsis has been observed. **(D)** Ploidy across primary-PDX-PDO trios (top). Genome-wide concordance score for primary-PDX, primary-PDO, and PDX-PDO pairs for 3 trios (PCSI_0590, PCSI_0624, and PCSI_0642) is indicated (bottom). The highest scoring sample for across tumour-PDX, tumour-PDO, and PDX-PDO pairs is highlighted in red.

Jaccard indices were computed to quantify overall similarity between the ploidy-adjusted copy number states of primary tumours and matching PDX, across all bases of the genome. Almost perfect concordance of copy number state was reflected genome-wide (concordance ≥ 0.8 in all primary-PDX pairs) [Figure 7B, Supplemental_Table_S8.pdf (A)]. However, copy number state was observed as highly variable across chromosomes for which clustered SV events were identified [Figure 7C]. Assessment of copy number variation in the liver metastasis set demonstrated comparable ploidy states between metastasis-PDX pairs [Figure 7A, Supplemental_Table_S7.pdf (B), Supplemental_Fig_S11.pdf], with almost perfect concordance of copy number state across all pairs (concordance rate ≥ 0.8) [Figure 7B, Supplemental_Table_S8.pdf (B), Supplemental_Fig_S12.pdf]. The MMR deficient sample (PCSI_0489) did not exhibit major disruptions in copy number across the genome [Figure 7A, Figure 6B, Supplemental_Fig_S12.pdf].

Copy number state was compared within tumour-PDX, tumour-PDO, and PDX-PDO matched pairs of trios [Figure 7D, Supplemental_Table_S7.pdf (C), Supplemental_Fig_S13.pdf]. As PCSI_0592 and PCSI_0602 were excluded due to ploidy estimation problems, only 3/5 trios were available for further analysis [Figure 7D, Supplemental_Fig_S13.pdf]. PDO samples exhibited similar ploidy to their matched primary tumour and PDX [Figure 7D, Supplemental_Table_S7.pdf (C)]. There was an overall concordance of ploidy between tumours, PDX, and PDO trios, with the exception of PCSI_0590, which exhibited disparate ploidy in the PDX as previously described [Supplemental_Fig_S14.pdf]. Pairwise-comparisons (tumour-PDX, tumour-PDO, and PDX-PDO) highlighted almost perfect concordance of copy number (concordance rate ≥ 0.8) in all the pairs [Figure 7].

## DISCUSSION

Critical evaluation of disease models is gaining importance across several cancer types that utilize these surrogates as preclinical tools for exploring tumourigenesis and drug testing. Previous investigations of PDAC provide a limited snapshot of donor-model comparisons, in terms of morphological, pathological, and mutational correlates [Supplemental_Fig_S15.pdf]. Our histologic analysis demonstrated that PDXs and PDOs retain the main characteristics of the tumour samples from which they were derived, in agreement with previous work **[4, 6, 8]**. It can be argued however, that sampling of multiple sites from donors and disease models can identify a spectrum of histologic and morphologic patterns that would render tumours and their matching models as dissimilar. Accordingly, donor-model comparisons of particularly heterogeneous tumours, including PDAC, will benefit from the advent of next-generation sequencing technologies that can encapsulate all facets of the disease at the molecular level. We present, to our knowledge, the first detailed exploration and quantitative assessment of whole-genome comparisons between human PDAC (both primary and metastasis) and matched model systems. We evaluated single base-pair genomic events (SSM), larger chromosomal changes across several bases or chromosomes (SV), and genome-wide changes (CNV) in tumour-PDX pairs and tumour-PDX-PDO trios.

In our PDAC cohort of paired tumour-PDX-PDO trios, the organoid samples are derived from the PDX, not the original patient donors. Given this experimental setup, the overall expectation is that the PDO would demonstrate greater fidelity with the PDXs. This was noted in our assessment of copy number variation across 3 trios: PDX-PDO pairs scored marginally more concordant than patient-PDO comparisons. Despite the variability observed across SV events, PDX-PDO pairs also demonstrated strong concordance, even in trios where patient-PDX and patient-PDO pairs are actually discordant (ex: PCSI_0602, PCSI_0642). Given our observation that PDX-PDO pairs score more concordant than patient-PDX or patient-PDO, this suggests that true PDO representation of the donor tumour would benefit from direct growth from the patient tumour itself.

Genomic comparisons between primary tissues and disease models are necessary for future studies that focus on gene-drug associations across model systems. Recent findings highlight that successful tumour engraftment of PDXs is associated with adverse clinico-pathological features and worse recurrence-free and overall survival; this variability in PDX growth suggests limited potential for systematic use of PDX tumours for real-time chemo-sensitivity testing **[20]**. Our approaches to score concordance and discordance of genomic events across donor-model pairs present quantifiable parameters of model fidelity, which can be used to support the utility of a given model for preclinical testing. For example, our analysis highlighted a discordant tumour-PDX pair (PCSI_0169) that was consistently difficult to decipher across both SSM and SV comparisons. One possible explanation for this discordance may be a mutational process that is only present in the PDX, due to a clone from the primary tumour growing as the main clone in the PDX. Further investigations would be needed to explore this phenomenon; however, this is a clear case where the PDX may not a good surrogate for preclinical testing of the donor tumour. Deciphering the extent of discordance for that PDX sample would not have been possible without scoring donor-model comparisons across genome-wide, single-base, and chromosome-centric scales.

At the single-base and gene-centric level, our findings emphasize that PDAC disease models successfully capture the same mutational patterns and driving events involved in PDAC tumourigenesis. We have scored genomic concordance of tumour-PDX pairs across different mutation types, and noted strong agreement between tumours (primary or metastasis) and their matched PDX. A more detailed evaluation also demonstrated retention of genetic features for 9 driver genes involved in PDAC tumourigenesis. As such, our study confirms and extends prior findings of fidelity of SSMs in tumours and their paired PDX **[4, 8, 21]**. We now demonstrate that this fidelity extends to PDOs as well. As the molecular landscape of PDAC is complex **[22]**, our observations that predicted deleterious functions of oncogenes and tumour suppressor genes remain faithfully represented in the PDXs and PDOs, provides further overall support for use of these models in precision medicine. PDOs are particularly attractive, as they can be readily established from small patient biopsies **[16]**.

On the genome-wide scale, quantitative scoring of copy number changes demonstrated high concordance between donor-tumour pairs for both PDX and PDO, most importantly for ploidy-scaled copy number events. While the majority of tumour-PDX pairs and trios analyzed demonstrated comparable ploidy, our analysis also captured conspicuous cases of whole genome polyploidization events that gave rise to tetraploid genomes in matched disease models (notably across PCSI_0590 and PCSI_0633). This deviation in ploidy underscores potential changes that arise when transferring portions of primary tumours to other disease model mediums, particularly across multiple passages. In our work, WGS profiling of PDX and PDOs was undertaken following third passage (P3) engraftment of tumours into mice. While the lack of profiling of earlier PDX passages prevents us from drawing conclusions about copy number aberrations (CNAs) in PDAC across multiple passages, one limitation includes the selection of subclones when different portions of the tissue are grown in future passages within PDXs and PDOs. Subclones may gain survival advantage growth in the disease models, and subsequently, tumour engraftment in PDXs may be confounded by underlying biological mechanisms that promote adaptation and growth of these tumours in a new environment **[20]**. Indeed, recent findings by Ben-David *et al*, based on observations from breast and hematopoietic cancers, suggest that clonal evolution of PDX occurs through directional selection of pre-existing clones **[23]**. Interestingly, their study emphasized quick genomic divergence and rapid CNA dynamics across the first few *in vivo* PDX passages, such that CNAs acquired through PDX passaging differ substantially from that in their parental tumours **[23]**. These revelations may explain the polyploidization events that we have observed in our PDAC PDXs, and the difficulty in attributing ploidy to some samples that were eventually excluded from the study (PCSI_0592 and PCSI_0602).

In comparison to SSM and CNV events, SV events captured the diverse range of PDAC heterogeneity across both tumours and disease models. Genome-wide scoring of SV concordance showed poor correlations between tumours and PDX, which promoted further investigation into chromosome-specific scoring methods that can efficiently depict genome-wide heterogeneity. We subsequently demonstrated that SV overall concordance (S^c^ score) fails to capture key instances of chromosome-specific concordance. Strikingly however, we identified cases of genomic concordance across chromosomes with clustered SV events that suggest chromothripsis. This phenomenon has not been previously described in PDAC disease models. Quantitative assessment of these chromosomes demonstrated concordance in tumour-PDX, tumour-PDO, and even PDX-PDO comparisons, compared with other chromosomes with fewer SV events. Our findings present a strong argument that, despite overwhelming evidence of structural heterogeneity between tumours and their matched models, major structural changes that occur in a tumour sample are effectively ‘transmitted’ to matched PDXs and PDOs. This is particularly reassuring in terms of translatability of these models when considering complex genomic events. There remain, however, other anomalous cases where clustered SV events in specific chromosomes within tumour samples have been ‘lost in translation’ in matched PDX and PDO (the most notable case is PCSI_0624, chromosome 17). Equally difficult to rationalize are cases where clustering of SV events have arisen in the matched PDX, but not in the source tumour (PCSI_0611, chromosome 22). While these events may be explained as a result of subclonal events or tumour selection, our limited sample size hinders investigations whether those events could be recurrent events in a larger PDAC series.

Our study presents a comparative analysis of patient tumours and disease models for both primary PDAC and metastasis. While our samples do not allow direct comparisons of matched primary and metastatic tumours, it is possible to draw conclusions about the overall genomic profile of the metastatic samples (and their matched PDX) in comparison with the primary series (and their matched PDX). Across all types of genomic events studied (SSM, SV, CNV), we observe that metastases demonstrate higher concordance with their matched PDX compared with primary-PDX pairs. All of the pairs demonstrate higher concordance levels (Jaccard index ≥ 0.6) for both SSM and CNV. SV heterogeneity across these pairs is less pronounced than those of primary tumours, producing improved concordance scores for SV events. One possible explanation for this behavior may reflect upon the tumour microenvironment of metastatic samples compared to primary tumours, as metastatic samples represent more stable, aggressive, and proliferative derivatives of the primary tumour. Interestingly, our cohort of metastasis-PDX pairs also includes a sample demonstrating DNA mismatch-repair (MMR) deficiency (PCSI_0489) due to a germline MLH1 mutation. Alterations in MMR genes lead to microsatellite instability (MSI), a genotype found infrequently in PDAC **[11, 24, 25]**. Our analysis of the MMR sample highlights a high mutation load, few SV events across chromosomes, and a stable DNA copy number across the genome; all of these observations are reflective of the genomic instability expected in tumour samples exhibiting MMR deficiency. Observing the same behavior in the matched PDX is reassuring, as it emphasizes that the PDX model succeeds in recapitulating much of the genomic behavior of these rare and striking PDAC cases.

This study has several potential limitations. Our limited sample size hampers investigations into subclonal events within individual samples (due to a lack of technical replicates). Due to a lack of sufficient biological replicates, our sample counts are too few to support comprehensive identification of recurrent genomic patterns across a larger cohort of PDXs and PDOs, including, for example, recurrently affected genes. Finally, we recognize that our chromosome-centric scoring of SV events between patient tumours and matching disease models relies on finite SV counts. This may present biased scoring for several chromosomes for which a very small number of events are tabulated, even though these events may span large genomic regions (for example, a deletion spanning several hundreds of bases). To overcome this, we delineated, as part of our calculations, the chromosomes whose SV events pass a minimum threshold (minimum of 5 events).

In summary, we have conducted detailed molecular dissection of WGS data to quantify concordance of genomic events between model systems and matched human PDAC. Our systematic comparison of tumours, PDX, and PDO highlight several genetic aberrations that are sample-specific, and which may not be shared across donor samples and matched models. Broadly, our findings indicate that PDX and PDO successfully recapitulate primary and metastatic PDAC at the level of SSMs and DNA copy number. However, disease model fidelity is not retained as well when assessing structural variation events, as evidenced by high variability observed at the level of tumour-PDX, tumour-PDO, and PDX-PDO comparisons. Strikingly however, clustering of SV events across particular chromosomes is retained when the tumours are implanted into their respective disease models. Collectively, our findings demonstrate that PDXs and PDOs serve as tractable and transplantable systems for probing the molecular properties of PDAC in terms of mutation and copy number changes, and for selective analysis of chromosome-specific structural variation events. We expect that our analytic pipeline may serve as a framework for future WGS research that compares donor samples and matched PDX and PDO.

## METHODS

A schematic overview of the biospecimens and analytic design is presented in Figure 1 and Supplemental_Fig_S1.

### Model system derivation

PDX were established by subcutaneous implantation of fresh surgically resected primary tumour tissue into immunodeficient mice **[26]**. All animal manipulations were approved by the University Health Network Animal Welfare Committee.

PDO models were generated by the Princess Margaret Living Biobank core facility using previously described protocols **[16]**. Briefly, fresh PDX tissue was cut into small pieces and dissociated to single cells or small clumps of cells using Liberase™ TH (Sigma Aldrich, Ontario, Canada). Dissociated cells were collected and embedded in growth factor-reduced Matrigel (Corning, New York, USA), which is overlaid with growth medium **[16]**

### Histopathology

Snap frozen tumour tissue 5mm^3^ or larger were obtained for each case and stored at −80°C. Each tissue was serially cryosectioned (10um thickness) at −20°C, fixed with 100% ethanol for up to 30 min and mounted onto PEN membrane 1.0 slides (Carl Zeiss MicroImaging, GmbH, Munich, Germany). All but one section was stained to visualize structures using a cresyl violet protocol that stains Nissl granules purple. Sections were rinsed in deionized water and stained in 1% cresyl violet solution (1% w/v in 50% ethanol, 50% deionized water) for 1 minute, dipped in 70% ethanol and dehydrated by dipping in absolute ethanol. Slides were left to dry for approximately 15 minutes before being stored at −80°C in air tight, aluminum foil wrapped slide boxes. The last section of each series was mounted onto an uncharged slide and stained using a modified haematoxylin and eosin (H&E) protocol. Sections were rinsed in deionized water, stained in Mayer’s haematoxylin (10 minutes), rinsed with deionized water, blued in Scott’s tap water, rinsed in deionized water, stained in aqueous eosin Y (1 minute), dipped in 3 changes of absolute ethanol, dehydrated in three changes of xylene (2 minutes each), prior to coverslipping. Additional details on the histochemistry and laser capture microdissection of xenograft samples have been previously described **[11]**.

### Laser-capture microdissection (LCM)

LCM was performed on the PDX as previously described **[11]**. Briefly, cresyl violet stained slides were brought to room temperature. Tumour cells from each cresyl violet section were microdissected using the PALM LMPC device (Carl Zeiss MicroImaging, GmbH, Munich, Germany). Tissue was collected in AdhesiveCap tubes (Carl Zeiss MicroImaging, GmbH, Munich, Germany) and stored at −80°C prior to extraction.

LCM of fresh frozen tissue samples from PDAC was performed on a Leica LMD 7000 instrument. Frozen tissue for tumour samples was maintained in vapor-phase liquid nitrogen and embedded in OCT cutting medium and sectioned in a cryotome into 8-μm thick sections. These sections were then mounted on PEN membrane slides (Leica) and lightly stained with hematoxylin to distinguish tumour epithelium from stroma. A pathologist (SF) marked tumour sections and LCM was performed on the same day according to manufacturer’s protocol on the Leica LMD7000 system. Microdissected tumour cells were collected by gravity into the caps of sterile, RNAse-free microcentrifuge tubes. Approximately 150,000 – 200,000 tumour cells were collected for each DNA extraction and stored at −80°C in Arcturus PicoPure Extraction Buffer.

### Whole genome sequencing

Paired-end cluster generation and sequencing was carried out using the Illumina HiSeq 2500 platform, on DNA isolated from fresh frozen tissue following LCM. Whole genome sequencing (WGS) of tumours, PDX, and PDO was performed with a minimum depth of ~30X per sample. Xenome (version 1.0.1) **[27]** was used to identify and filter mouse content. Non-mouse DNA reads from primaries, PDX and PDO were aligned to the human reference genome hg19 using the Burrows-Wheeler Aligner (BWA, version 0.6.2) **[28]** with default parameters. Picard (version 1.90) (**http://broadinstitute.github.io/picard**) was used to sort, merge, and mark duplicates from multiple lanes of the same sample, followed by the Genome Analysis Toolkit (GATK, version 1.3.16) **[29, 30]** to improve alignment accuracy.

### Simple Somatic Mutations (SSM)

Germline single nucleotide variants (SNV) were called using the Genome Analysis Tool Kit (GATK, version 1.3.16), using best practice guidelines made available by the Broad Institute. Briefly, data were locally realigned around indels and the base quality values were recalibrated prior to variant calling using the Unified Genotyper. This was followed by filtering using the VariantFiltration module, and subsequent classification of germline variants as those mutations which have a QUAL score greater than 50 in the normal sample. Both tumour and matched normal samples were processed simultaneously.

Strelka (version 1.0.7) **[31]** and MuTect (version 1.14) **[32]** were used to call somatic SNVs, with default parameters. Indels were also identified using Strelka. SNVs were selected based on the intersection of ‘Tier 1 SNVs’from Strelka and ‘PASS’ filter variants from MuTect. Potential false positives caused by unfiltered mouse DNA were filtered using a blacklist of SNVs and INDELS generated by aligning model mouse DNA to hg19. Germline and somatic SSM were annotated using dbSNP 142 **[33]**, COSMIC (version 54) **[34]**, and ANNOVAR (version 2013-06-21) **[35]** to predict coding consequences of SNVs and indels. Functional consequences of mutations were predicted using Oncotator (version 1.5.3.0) **[36]**.

Parsing of VCF files containing the filtered calls was conducted using the *vcfR* (version 1.4.0) **[37]** and *VariantAnnotation* (version 1.18.7) **[38]** packages in R. Measurements of read depth of the variant and reference alleles were extracted to plot the frequency of reads carrying the variant allele. Assessment of mutation patterns for PDAC driver genes across all samples was performed by parsing the output generated by Oncotator using the *GenVisR* package (version 1.0.4) **[39]**.

Concordance of mutation calls between a tumour and its matching PDX was calculated using the Jaccard Index. For each mutation category annotated by Oncotator, the Jaccard Index for that mutation type (J_α_) was calculated as follows:

where

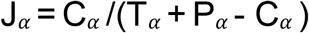

*α* = Oncotator mutation type (ex: lincRNA, missense mutation)
T_*α*_ = Number of variant calls in the tumour sample annotated with mutation *α*
P*_α_* = Number of variant calls in the PDX annotated with mutation *α*
C*_α_* = Number of variant calls in both the tumour and PDX that are annotated with mutation *α*

### Structural Variation

Structural variations (SV) were called using CREST (version alpha) **[40]** and DELLY (version 0.5.5) **[41]** with default parameters, and high-confidence SVs subsequently filtered. For high-confidence SV calls observed in at least one sample of a tumour-PDX pair, we manually reviewed whether the same variant was observed in the matched sample with lower frequency, and added those ‘rescued’ variants to the filtered list. The union of the filtered calls and the rescued calls was used for all downstream analysis.

Genome-wide structural changes across all the four categories of structural variation (DEL = deletion, INV = inversion, DUP =duplication, and TRA = translocation) in tumour-PDX pairs were rendered using the *RCircos* library (version 1.2.0) **[42]**. Quantification of structural variation events in all PDAC and liver metastasis tumour-PDX pairs was calculated using functions from the *RCircos* (version 1.2.0) **[42]**, *GenomicRanges* (version 1.24.3) **[43]**, *rtracklayer* (version 1.32.2) **[44]** and *PharmacoGx* (version 1.1.6) **[45]** packages in R.

Assessment of concordance and discordance among SV events was conducted for each chromosome individually across tumour-PDX pairs. We identified chromosomes with ≥ 5 SV events in both tumours and their matching PDX. To count SV events per chromosome, intra-chromosomal events (deletions, inversions, duplications) were assigned a score of ‘1’ for their respective chromosomes, while a score of 0.5 was assigned to each of the chromosomes involved in a translocation event. Instances of discordance (where one sample of a pair had ≥ 10 SV events different from the other sample) were also noted.

Concordance of structural variation events for a tumour and its matching PDX was quantified using the Jaccard Index. Jaccard indices of a tumour-PDX pair were calculated individually for each chromosome (J_β_), across all SV events as follows:

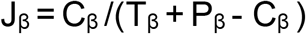

β = chromosome (autosomes in addition to chrX and chrY)
T_β_ = Number of SV calls in the tumour in chromosome β
P_β_ = Number of SV calls in the PDX in chromosome β
C_β_ = Number of SV calls in both the tumour and the PDX in chromosome β

We also generated an overall concordance (S^c^) score to summarize the agreement between tumour-PDX pairs. The score determines the ratio of ‘positive’ chromosomes across a tumour-PDX pair that have a Jaccard index ≥ 0.6. The score is calculated as follows:

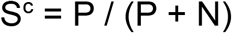

where

P = number of chromosomes with J_β_ ≥ 0.6
N = number of chromosomes with J_β_ < 0.6

In the case of trios (tumour-PDX-PDO), SV concordance was quantified by splitting the trio into pairwise calculations of tumour-PDX, tumour-PDO, and PDX-PDO, and then calculating J_β_ and S^c^, as described previously.

### Copy number variation

Copy number segments were obtained using CELLULOID (version 0.11.2) to estimate gene copy number and tumour ploidy from WGS **[12]**. Unless otherwise specified, copy number segments and parameters were extracted for the first solution (solution1) of the CELLULOID proposed solutions, for each sample.

Concordance of copy number state was calculated by first identifying overlapping genomic loci in tumour-PDX pairs using bedtools (version 2.24.0) **[46]**. To consider the copy number state relative to ploidy, the *imean* scores generated by CELLULOID (which represents average integer copy-number) for the genomic loci were rescaled by the ploidy of the respective samples. Genomic loci in tumour-PDX pairs were considered identical if they shared a copy number state with a difference ≤ 0.25. Genome-wide concordance scores (G) were calculated across all bases of a tumour-PDX pair as follows:

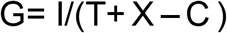

T = Number of genomic bases in the tumour with a defined copy number (imean-rescaled value)
X = Number of genomic basis in the PDX with a defined copy number (imean-rescaled value)
C = Number of genomic basis in either the tumour or PDX with a defined copy number (imean-rescaled value)
I = Subset of C, where the absolute value of the difference between the imean score in the tumour and PDX is ≤ 0.25

In the case of trios (tumour-PDX-PDO), CN concordance was quantified by splitting the trio into pairwise calculations of tumour-PDX, tumour-PDO, and PDX-PDO, as above. Plots of overlapping copy number states were drawn using the *copynumber* package (version 1.12.0) **[47]**.

### Ranking concordance between models

Quantitative comparison of SSM, SV, and CNV events between tumours and their corresponding PDX and PDO samples were developed by calculating Jaccard indices (J_α_, J_β_) or concordance scores (S^c^, G), as described previously. These scores follow a scale from 0 to 1. To facilitate comparison of different models across these different metrics, these scores can be qualitatively described in terms of overall concordance as follows: 0–0.2: strongly discordant; 0.2–0.4: discordant; 0.4–0.6: moderate; 0.6–0.8: strongly concordant; 0.8–1.0: almost perfectly concordant.

## DATA ACCESS

The datasets analyzed during the current study are available in the European Genome-phenome Archive (EGA), accession code EGAS00001002597. Comparison of SSM, SV, and CNV events between tumours and their corresponding PDX and PDO sample was conducted using R (version 3.3.1) **[48]**. All software dependencies are available on Bioconductor (BioC) or the Comprehensive Repository R Archive Network (CRAN), and have been listed throughout the methods as applicable. The code and associated tutorial describing how to run the analysis pipeline are publicly available on Github (github.com/DGendoo/PDACDiseaseModels).

## AUTHOR CONTRIBUTIONS

D.M.A.G, S.G, and B.H-K conceived the initial design of the study.

D.M.A.G designed and developed the analysis, wrote the code, performed the analysis and interpreted the results.

R.E.D and A.Z provided processed WGS data files (TSV and VCF files) for SSM and SV analysis and interpreted the results under the supervision of L.D.S and S.G.

G.H.J revised the statistical analysis.

M.L provided CELLULOID data files for CNV analysis.

S.F performed histo-pathological examination and diagnosis of the tissue samples.

D.C, and I.M.L conducted LCM of tissues under supervision of J.M.S.B.

N.R., E.I. established primary PDX and all PDO models under supervision of M-S.T. P-J.C. established metastasis PDX models under supervision of D.H.

N.D and D.H provided samples pertaining to the metastasis-PDX cohort.

J.M.W managed and supervised sample acquisition, annotation and data collection for PanCuRx.

D.M.A.G, R.E.D, N.R, J.M.W, S.G, and B.H-K reviewed and edited the manuscript.

S.G, and B.H-K supervised the study.

D.M.A.G, S.G, and B.H-K wrote the manuscript.

All authors read and approved the final manuscript.

## ACKNOWLEDGEMENTS

We thank Faiyaz Notta and Ashton Connor for comments and suggestions on the manuscript and analysis pipeline. We also acknowledge the technical contributions of the following individuals for their roles in Production Sequencing and Genomic Sequencing Informatics at the Ontario Institute for Cancer Research: Jenna Eagles, Jeremy Johns, Xuemei Luo, Faridah Mbabaali, Jessica Miller, Tanya Mohanta, Danielle Pasternack, and Morgan Taschuk. This study was conducted with the support of the Ontario Institute for Cancer Research (OICR, PanCuRx Translational Research Initiative) through funding provided by the Government of Ontario (Ministry of Research, Innovation, and Science), and a charitable donation from the Canadian Friends of the Hebrew University (Alex U. Soyka). D.M.A.G was supported by the PanCuRx Translational Research Initiative at the OICR. B.H.K was supported by the Gattuso Slaight Personalized Cancer Medicine Fund at Princess Margaret Cancer Centre, the Canadian Institutes of Health Research, the Natural Sciences and Engineering Research Council of Canada, and the Ministry of Economic Development and Innovation/Ministry of Research & Innovation of Ontario (Canada). N.R., M.S.T. and the establishment of PDO models are partially supported by fund from the Princess Margaret Cancer Foundation.

## SUPPLEMENTAL FIGURES

**Supplemental_Fig_S1.pdf: Schematic overview of study analysis**.

**Supplemental_Figure_S2.pdf: Immunohistochemistry staining (H&E, CK19) for PDX and PDO samples of the trios**.

**Supplemental_Fig_S3.pdf: Frequency of reads carrying the variant allele for primary-PDX pairs**. Oncogenes, tumour suppressors, and genes involved pathways of PDAC tumourigenesis are plotted.

**Supplemental_Fig_S4.pdf: Frequency of reads carrying the variant allele for metastasis-PDX pairs**. Oncogenes, tumour suppressors, and genes involved in pathways of PDAC tumourigenesis are plotted.

**Supplemental_Fig_S5.pdf: Frequency of reads carrying the variant allele for the trios**. The trio is split into primary-PDX (top), primary-PDO, and PDX-PDO (bottom) pairs. Oncogenes, tumour suppressors, and genes involved in pathways of PDAC tumourigenesis are plotted.

**Supplemental_Fig_S6.pdf: Circos plots of SV events across the genomes of each primary-PDX pair (n=10 samples)**. Each type of SV event is color-coded with a similar color between tumours and matching PDX. For each SV type, tumours are annotated on the outer rings of the circos plot and the matching PDX on the inner rings. SV events are colored as follows: deletions (red), inversions (green), and duplications (blue). Translocation events between chromosomes are also depicted (center).

**Supplemental_Fig_S7.pdf: Circos plots of SV events across the genomes of each metastasis-PDX pair (n=6 samples)**. Each type of SV event is color-coded with a similar color between tumours and matching PDX. For each SV type, tumours are annotated on the outer rings of the circos plot and the matching PDX on the inner rings. SV events are colored as follows: deletions (red), inversions (green), and duplications (blue). Translocation events between chromosomes are also depicted (center).

**Supplemental_Fig_S8.pdf: Circos plots of SV events across the genomes of each primary-PDX-PDO trio (n=5 samples)**. Each type of SV (deletion, inversion, duplication, and translocation) is represented as one circos plot, with 3 rings indicating tumour (outer), PDX (middle), and PDO (inner). SV events are colored as follows: deletions (red), inversions (green), and duplications (blue). Translocation events between chromosomes are also depicted (center).

**Supplemental_Fig_S9.pdf: Copy number profile across 8 matched primary-PDX pairs, as rendered by CELLULOID**.

**Supplemental_Fig_S10.pdf: Copy number across primary-PDX pairs for 8 samples**. For each pair, the copy number state of the primary tumour (magenta) and matching PDX (black) is plotted across all chromosomes. A detailed panel also shows the copy number of the tumour and PDX across each chromosome.

**Supplemental_Fig_S11.pdf: Copy number profile across 6 matched metastasis-PDX pairs, as rendered by CELLULOID**.

**Supplemental_Fig_S12.pdf: Copy number across metastasis-PDX pairs for 6 samples**. For each pair, the copy number state of the metastatic tumour (magenta) and matching PDX (black) is plotted across all chromosomes. A detailed panel also shows the copy number of the tumour and PDX across each chromosome.

**Supplemental_Fig_S13.pdf: Copy number profile across 3 primary-PDX-PDO trios, as rendered by CELLULOID**. Complete CELLULOID solutions (solutions 1-5) are also provided for each of the samples which have been excluded from the analysis (PCSI_0592 and PCSI_0602).

**Supplemental_Fig_S14.pdf: Copy number across 3 primary-PDX-PDO trios**. For each trio, the copy number of the tumour (black), matched PDX (red), and matching PDO (blue) is plotted across the genome. A detailed panel also shows the copy number of the tumour and PDX across each chromosome.

**Supplemental_Fig_S15.pdf: Review of the current literature comparing PDAC tumours and disease models**.

## SUPPLEMENTAL TABLES

**Supplemental_Table_S1.xls: Meta-data of the samples studied**.

**Supplemental_Table_S2.xls: IHC (H&E and CK19) observations pertaining to PCSI samples**.

**Supplemental_Table_S3.xlsx: Total number of simple somatic mutation (SSM) calls across the cohorts**. SSM calls across **(A)** primary tumours and matched PDX, **(B)** metastatic tumours and matched PDX, and **(C)** primary-PDX-PDO trios. Jaccard index for each tumour-PDX pair for mutation types are also shown for **(D)** primary tumours and matched PDX and **(E)** metastatic tumours and matched PDX.

**Supplemental_Table_S4.xlsx: Total number of structural variants (SV) calls across the cohorts**. SV across **(A)** primary tumours and matched PDX, **(B)** metastatic tumours and matched PDX, and **(C)** primary-PDX-PDO trios. For each sample, the total number of deletions (DEL), duplications (DUP), inversions (INV), and translocations (TRA) is indicated.

**Supplemental_Table_S5.xlsx: Total number of structural variation events (SV) observed in each chromosome**. Total events are indicated across **(A)** primary tumours and matched PDX, **(B)** metastatic tumours and matched PDX, and **(C)** primary-PDX-PDO trios. Deletions, duplications, and inversion events that occur in a chromosome were assigned a value of 1 prior to their summation, and translocation events assigned a value of 0.5.

**Supplemental_Table_S6.xlsx: Chromosome-specific Jaccard indices**. The indices are indicated across **(A)** primary tumours and matched PDX, **(B)** metastatic tumours and matched PDX, and **(C)** primary-PDX-PDO trios.

**Supplemental_Table_S7.xlsx: Celluloid metrics for each of the samples**. Metrics are shown across **(A)** primary tumours and matched PDX, **(B)** metastatic tumours and matched PDX, and **(C)** primary-PDX-PDO trios. For each sample, the percentage of normal content (N), percentages of tumour content (T1), and the ploidy of the sample is indicated.

**Supplemental_Table_S8.xlsx: Genome-wide concordance score for copy number**. Concordance is computed across **(A)** 8 primary-PDX pairs and **(B)** 6 metastasis-PDX pairs.

